# Saccade-related modulation of oscillatory activity in primary auditory cortex

**DOI:** 10.1101/2022.05.10.491383

**Authors:** Annamaria Barczak, Monica Noelle O’Connell, Tammy McGinnis, Kieran Mackin, Charles E. Schroeder, Peter Lakatos

## Abstract

The auditory and visual sensory systems are both used by the brain to obtain and organize information from our external environment, yet there are fundamental differences between these two systems. Visual information is acquired using systematic patterns of fixations and saccades, which are controlled by internal motor commands. Sensory input occurs in volleys that are tied to the timing of saccades. In contrast, the auditory system does not use such an overt motor sampling routine so the relationship between sensory input timing and motor activity is less clear. Previous studies of primary visual cortex (V1) in nonhuman primates (NHP) have shown that there is a cyclical modulation of excitability tied to the eye movement cycle and suggests that this excitability modulation stems from the phase reset of neuronal oscillations. We hypothesized that if saccades provide a supramodal temporal context for environmental information then we should also see saccade-related modulation of oscillatory activity in primary auditory cortex (A1) as NHPs shift their gaze around their surroundings. We used linear array multielectrodes to record cortical laminar neuroelectric activity profiles while subjects sat in a dark or dimly lit and silent chamber. Analysis of oscillatory activity in A1 suggests that saccades lead to a phase reset of neuronal oscillations in A1. Saccade-related phase reset of delta oscillations were observed across all layers while theta effects occurred primarily in extragranular layers. Although less frequent, alpha oscillations also showed saccade-related phase reset within the extragranular layers. Our results confirm that saccades provide a supramodal temporal context for the influx of sensory information into A1 and highlight the importance of considering the effects of eye position on auditory processing.

**Significance Statement:** Using laminar multielectrodes, the current study examined saccade-related neuronal activity during resting state while NHPs sat in a dark or dimly lit room. Our results confirm that saccade-related modulation of delta band oscillatory activity occurs across all layers of A1. Interestingly, our data also show a saccade-related phase reset of theta and alpha bands that preferentially occurs in extragranular layers. These results confirm that saccades provide a supramodal temporal context for the influx of environmental information into A1 and emphasizes the importance of considering eye position when examining auditory processing.

## INTRODUCTION

Through the process of active sensing, sensory information from the external world is sought out using internally controlled motor sampling routines
(Kleinfeld et al., 2006; Schroeder et al., 2010; Ahissar and Arieli, 2012). The visual and auditory sensory systems both obtain and organize outside information. However, in human and non-human primates (NHPs), sensory exploration in the visual and auditory modalities differ greatly. During visual active sensing, information is mainly acquired using systematic patterns of fixations and saccades. Visual input occurs in volleys “caused” by saccades and is strictly tied to saccade timing. While auditory active sensing utilizes head movements to maximize binaural cues (Thurlow et al., 1967), these movements do not initiate volleys of auditory input so the relationship between sensory input timing and motor activity is less clear.

Ongoing neuronal oscillations reflect alternating states of neuronal ensemble excitability (Buzsáki, 2006) and oscillations within certain frequency bands can be recruited by sensory inputs to enhance stimulus processing (Buzsáki and Draguhn, 2004; Lakatos et al., 2005). These oscillations in motor (Lalo et al., 2007) and visual (Rajkai et al., 2008; Ito et al., 2013; Barczak et al., 2019) cortices can also be enlisted by the motor system during active sensing. The quasi-rhythmic organization of incoming visual information by saccades allows for an ideal opportunity to align with ongoing oscillations and lead to more effective stimulus processing. These effects were previously demonstrated in NHPs whereby rhythmically occurring saccades were shown to result in a biphasic modulation of excitability: saccade-related suppression followed by fixation-related enhancement in V1 (Rajkai et al., 2008; McFarland et al., 2015; Barczak et al., 2019). This saccade-related organization of visual cortical neuronal activity in NHPs has been observed both during free-viewing (Ito et al., 2013) and in the dark (Rajkai et al., 2008; Barczak et al., 2019). The latter studies suggested that phase reset of neuronal oscillations across multiple discrete frequency bands might underlie the biphasic modulation of neuronal excitability, and since these effects were observed with a lack of visual input (i.e. dark environment), the authors concluded that they are likely due to a nonretinal, motor-related component of the eye movement signal.

While saccade-related effects across the visual pathway have been widely observed, the degree to which saccade-related, possibly motor-initiated, effects on non-visual sensory areas has been less explored. Importantly, while saccade-related modulation of primary auditory cortex (A1) could be due to motor signals related to the saccade (e.g. corollary discharge signal), given that other sensory modalities have been shown to modulate activity at the level of primary auditory cortex (Lakatos et al., 2007; Kayser et al., 2008; Musacchia and Schroeder, 2009), it is also possible that observed effects could be due to multisensory interactions from the visual input associated with each saccade end (i.e. fixation onset). Recently, saccade-related eardrum motion has been observed in the absence of auditory stimuli and this motion appeared to have oscillatory characteristics (Gruters et al., 2018). Additionally, saccade-related entrainment and modulation of excitability has been shown to occur in A1 of NHPs (O’Connell et al., 2020). Although perceptual saccadic suppression of auditory stimuli presented during saccades has not yet been demonstrated (Harris and Lieberman, 1996), saccades have been shown to influence the perception of auditory stimulus localization (Krüger et al., 2016).

Building on our previous results (O’Connell et al, 2020), the main goal of the current study was to further examine saccade-related entrainment in A1. Linear array multielectrodes were used to observe neuroelectric activity across all layers of A1 in macaque monkeys, while eye movements were monitored as they sat passively in a dark or dimly lit room. By separating modulation related to saccade onset vs. offset, we aimed to differentiate between effects due to motor (predictive) signals and those related to visual input (reactive). We found significant saccade-related modulation of activity in A1 where the main effect was a phase reset (Makeig et al., 2002; Rizzuto et al., 2003; Sauseng et al., 2007; Arnal and Giraud, 2012) of neuronal oscillations in the delta, theta, and alpha frequency bands. Phase reset of oscillations in the delta and theta frequency bands could reliably be linked to both saccade onset and offset, while alpha phase reset also occurred – albeit much less frequently. Modulation of delta was observed across all cortical layers, while theta and alpha effects occurred more often in extragranular layers. The overall results suggest that saccade-related, “motorsensory” modulation is supramodal and is therefore an ideal mechanism to coordinate activity across multiple brain regions during active visual sampling of the environment.

## METHODS

### Subjects

All procedures were approved in advance by the Animal Care and Use Committee of the Nathan Kline Institute. Eye position and neuroelectric activity were recorded from 3 macaques (M. mulatta, 2 females/1 male, 4.2-9.0 kg) obtained from an approved source. Data from a total of 62 recording sites gathered across 52 experimental sessions were used (33, 23, and 6 sites from 27, 19, and 6 sessions with macaques R, B, and M, respectively). During 10 of the experimental sessions, two electrodes were each positioned in A1 within different hemispheres.

### Surgical & Behavioral Preparation

Surgical preparation of subjects for awake intracortical recordings was performed under general anesthesia using aseptic techniques. To permit access to the brain and orthogonal placement of electrodes in auditory cortex, Polyetheretherketone (PEEK; Rogue Research Inc.) recording chambers were positioned and secured normal to the cortical surface of the superior temporal plane using ceramic screws. Known stereotaxic coordinates and the results from preoperative structural MRIs for each NHP were used to guide this surgical placement. Since head immobilization was required for acute electrode recordings and eye tracking, a PEEK headpost was secured to the skull with ceramic screws and embedded in dental acrylic to allow for painless head restraint. Each NHP was given a minimum of 6 weeks for post-operative recovery before behavioral training and data collection began.

NHPs were adapted to a custom fitted primate chair and to the recording chamber. After the post-surgical recovery period, NHPs were acclimated to handling and were brought to the laboratory 2 to 3 times per week and trained to allow painless head restraint.

### Electrophysiological Recording

All recordings were performed in a dark or dimly lit, electrically shielded and sound attenuated chamber. The broadband neuroelectric signal was recorded using the Alpha Omega SnR (Alpharetta, GA) system with a 44 kHz sampling rate. Linear array multielectrodes (23 equally spaced contacts with 100 μm intercontact spacing; Fig. 1A) were acutely positioned to sample neuroelectric activity from all cortical layers of A1 simultaneously. To aid electrode positioning, broadband noise bursts (BBN) were presented (1.6 Hz repetition rate) and response profiles were created and used as a guide.

**Figure 1.**
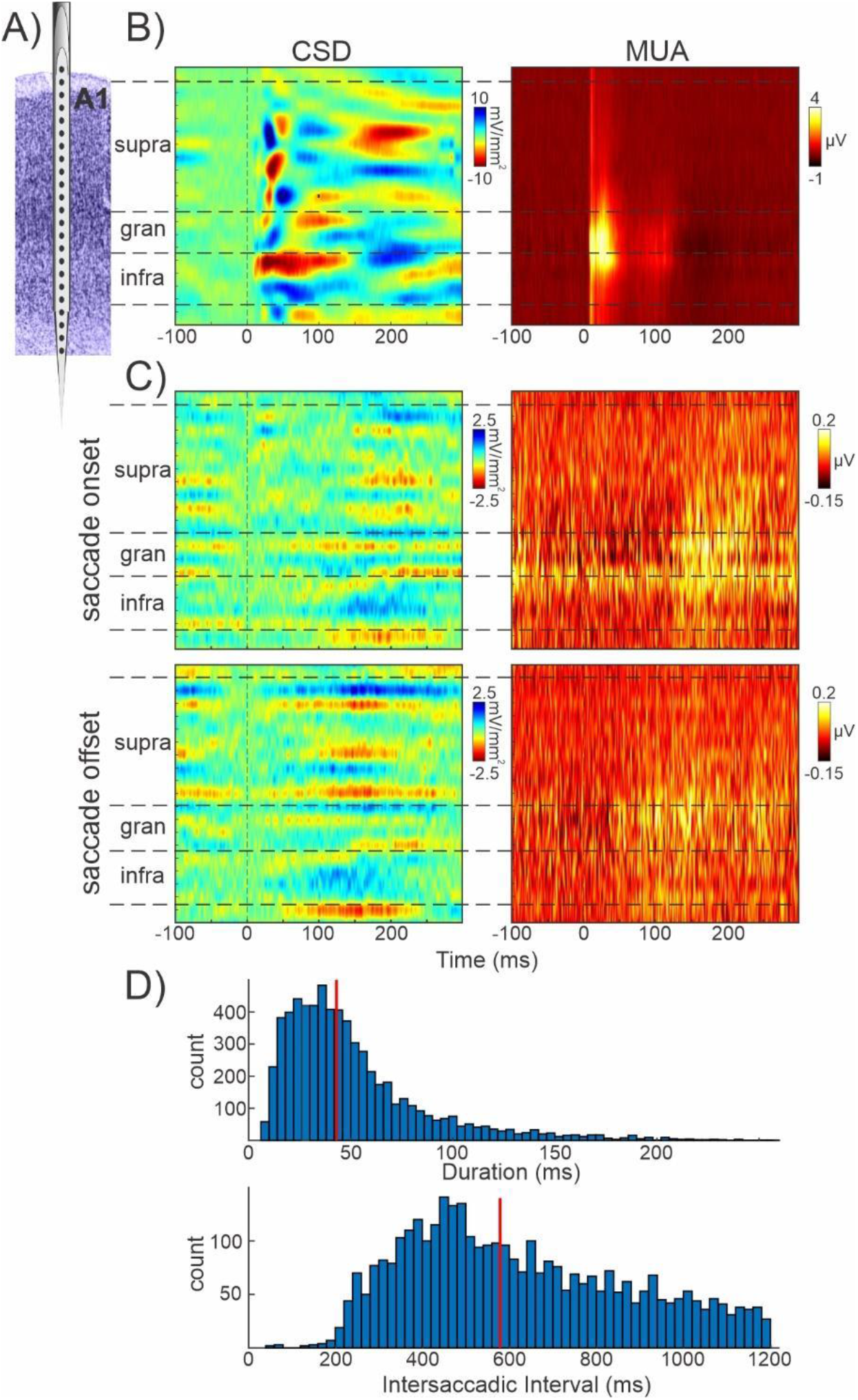
Saccade-related modulation of CSD and MUA in A1. **(A)** Schematic of a laminar multielectrode spanning all cortical layers of A1. **(B)** Representative CSD (left) and MUA (right) response profiles recorded from A1, aligned to the onset of a broadband noise burst. Horizontal dashed lines show approximate boundaries between supragranular, granular, and infragranular layers. **(C)** Representative CSD and MUA profiles from the same A1 site as in panel B, aligned to saccade onset (top) and saccade offset (bottom). **(D)** Distribution of individual saccade durations (top) and intersaccadic intervals (bottom) of all 6468 saccades detected and verified through manual inspection. Red vertical lines indicate the median saccade duration and median intersaccadic interval. Intersaccadic intervals greater than 1.2 seconds were excluded from this figure.

Data analyzed in the current manuscript were collected while NHPs sat in a dimly lit or dark recording chamber. For a period of 3-5 minutes during recording, no stimuli were presented, and no rewards were given. These periods are referred to as a resting state condition. All offline analyses were conducted using custom-written and industry provided scripts and functions within Matlab (MathWorks, Inc, Massachusetts).

### Eye Tracking & Saccade Detection

Although their head was immobilized, NHPs were free to move their eyes around the dark chamber without restrictions. Eye position was monitored using the EyeLink 1000 system (SR Research Ltd., Mississauga, Canada) with the sampling rate set to either 500 or 1000 Hz. Analog eye position information from EyeLink was recorded at 2750 Hz concurrent to electrophysiological recordings using the Alpha Omega SnR system. Before any eye movement related analyses, the resolution rate of all analog eye movement data was set to 1000 Hz. Saccade onsets and offsets were determined using a nonparametric algorithm, Cluster Fix (König and Buffalo, 2014), which relies on k-means cluster analysis to determine event timing without having to set predefined thresholds. Manual verification of the detected saccades and rejection of possible artifacts was completed by two lab members. Across 62 experimental sessions, a total of 22,328 saccades were detected with the Cluster Fix algorithm (median number per experiment was 359). From this total, 6468 saccades were manually verified. Only experimental sessions with at least 40 manually verified saccades were included in these analyses. The median number of verified saccades across all 62 experiments was 93.

While the operational definition of fixations and saccades can vary across studies (Hessels et al., 2018), the current set of analyses focuses on saccade onset and offset as determined by the above algorithm. Manual inspection ensured that detected saccades showed a sharp initial change in X and/or Y eye position. Although we did not empirically evaluate velocity changes in X and Y eye position, visual inspection suggested that the timing of the saccade offsets determined by the algorithm did not always coincide with a stable X and Y eye position but instead often coincided with a decrease in the velocity of X and/or Y position changes. The current study did not include verification that periods immediately after saccade offsets could be characteristically defined as fixations (extended periods of stable X and Y eye position). Therefore, results related to saccade offset may not directly be related to those associated with fixation onset.

### Data Analysis, Signal Processing and Statistical Analyses

Neuroelectric signals were separated offline with zero phase shift Butterworth bandpass filters into field potential (0.1-300 Hz) and multiunit activity (MUA) (300-5000 Hz). One-dimensional CSD profiles were calculated from the LFP profiles using a three-point formula for the calculation of the second spatial derivative of voltage (Nicholson and Freeman, 1975). To compare auditory and saccade on/offset related response profiles, CSD and MUA activity were aligned to BBN stimulus onset and saccade on/offset (Fig. 1).

For the analysis of neuronal oscillatory activity, instantaneous oscillatory amplitude and phase were extracted from CSD signals using wavelet decomposition with the Morlet wavelet (omega=6) on 82 scales from 0.76 Hz to 106.48 Hz. Wavelet transformed data were epoched (segmented) according to the timing of saccade events (saccade onset and offset). For each frequency and channel, oscillatory amplitude was averaged across these epochs. Intertrial coherence (ITC) was calculated from oscillatory phases related to saccade events for each frequency and channel. ITC values indicated how clustered oscillatory phases were around a mean with 0 corresponding to completely random and 1 corresponding to perfectly aligned phases.

To evaluate saccade-related effects on oscillatory activity in A1 (Fig. 2), time-frequency ITC was averaged across all channels spanning A1 and all recordings (Fig. 2A). For the frequency specific averages, ITC and oscillatory amplitude within delta, theta, and alpha frequency bands were averaged across all channels (from top to bottom of cortex) and experiments (Fig. 2B, left). Peak timings for these averages were defined as the time points of the max ITC occurring within 0-300 ms relative to saccade onset and offset. These peak times were used for the statistical comparisons shown in the boxplots (Fig. 2B, right). For each individual experiment, “base” ITC was the average from −300 to 300 ms around saccade on/offset while “peak” ITC was the average across a 100 ms period that centered on the timing of the peak ITC (vertical dashed line in Fig. 2B). Oscillatory amplitude averages (Fig. 2C, left) for each experiment were normalized by dividing by the mean amplitude from −300 to 300 ms around saccade onset and offset. In Fig. 2C, boxplots on the right show the average normalized amplitude for each experiment for the same two “base” (−300 to 300 ms) and “peak” (100 ms centered on the ITC peak) time periods. For the examination of layer specific saccade-related effects (Fig. 3), ITC and oscillatory amplitude were averaged within delta, theta, and alpha bands and across channels within the supragranular, granular, and infragranular layers of A1. Averages and peak ITC timing were determined for layer specific ITC as described above.

**Figure 2.**
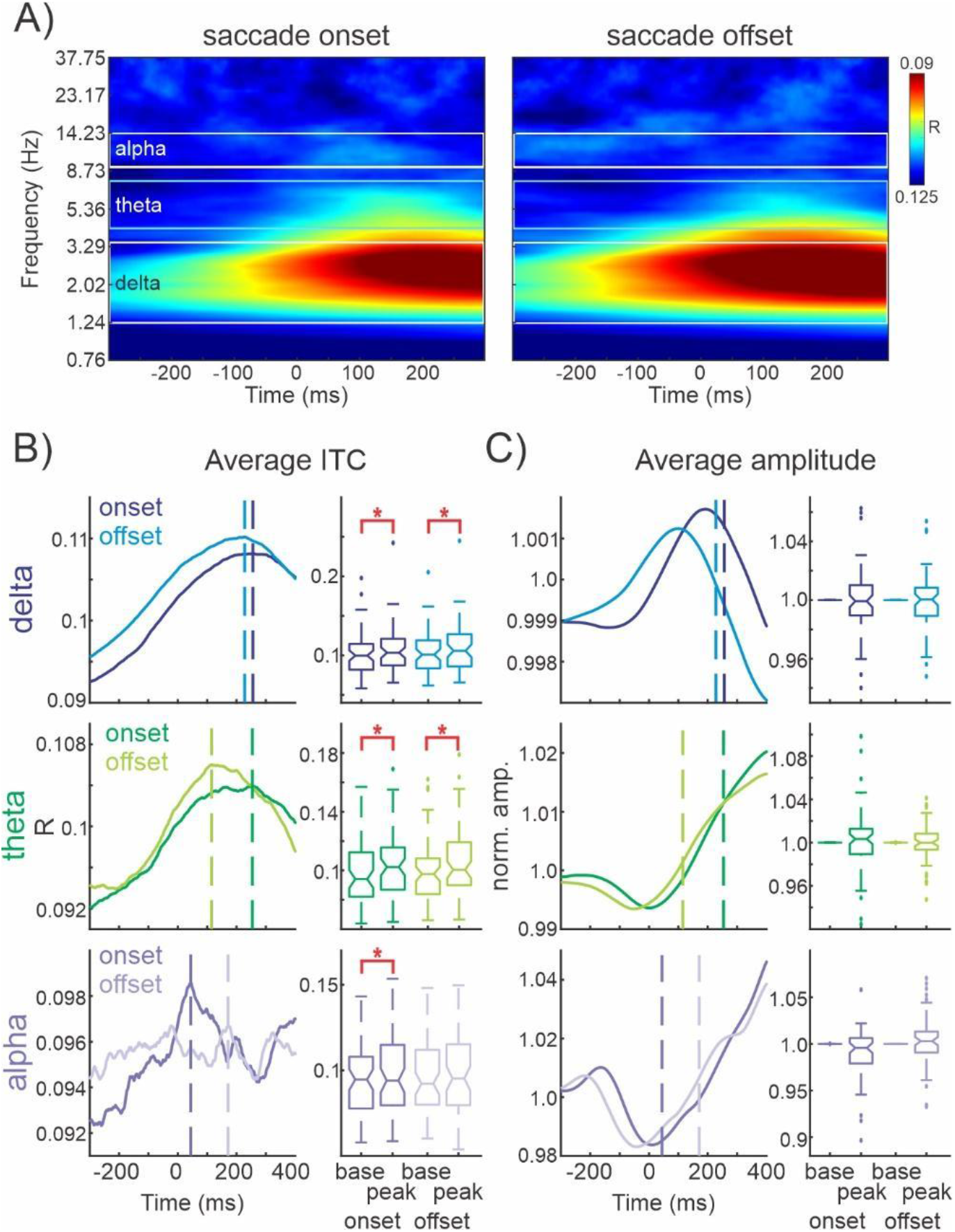
Saccade-related effects on oscillatory activity in A1. **(A)** Time-frequency ITC averaged across all channels (median # channels for each A1 site=18) and recordings (n=62) aligned to saccade onset (left) and saccade offset (right). Frequency bands of interest are outlined in white (delta/alpha) or cyan (theta) horizontal boundaries. **(B)** ITC aligned to saccade onset and offset were averaged within frequency bands of interest and across all cortical channels within each experiment. Average delta, theta and alpha ITC is shown on the left (top, middle, bottom, respectively). Boxplots on the right show average baseline ITC (base; from −300:300ms) and the average ITC 50 ms before and after the timing of the peak average ITC indicated by vertical dashed lines on left panels across each of the 62 experiments (peak). Red brackets indicate significant differences based on Wilcoxon signed rank tests (Bonferroni corrected). Delta saccade onset p=1.3852×10-^6^, saccade offset p=2.3248×10-^5^; theta onset p=0.0001, offset p=2.5024×10-^6^; alpha onset p=0.0013, offset p=0.7450. **(C)** Saccade onset and offset-related oscillatory amplitudes were averaged within frequency bands of interest and across all cortical channels within each experiment. General organization is the same as (B). Boxplots on the right show normalized baseline amplitudes (base; from −300:300ms) and the average amplitude 50 ms before and after the timing of the peak average ITC as determined in (B) (note: vertical dashed lines in B and C are the same). Some outliers were excluded from the boxplots for visual purposes but were included in all statistical evaluations. Number of outliers excluded (# onset/# offset) was delta: 2/2, theta: 2/1, alpha: 3/1. There were no significant amplitude differences based on Wilcoxon signed rank tests (Bonferroni corrected). Delta saccade onset p=0.9692, saccade offset p=0.5869; theta onset p=0.3639, offset p=0.7444; alpha onset p=0.0170, offset p=0.1256.

**Figure 3.**
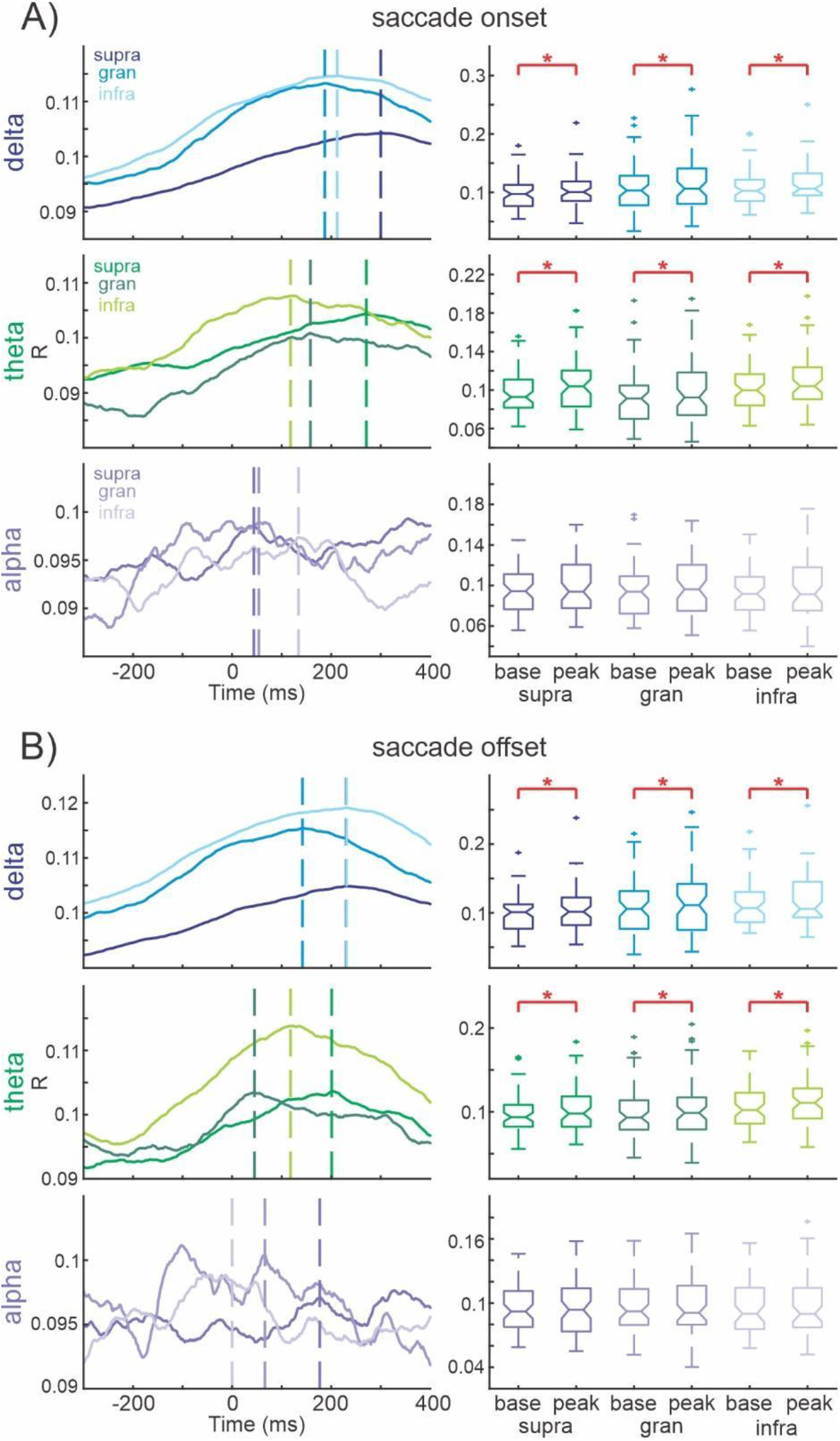
Layer specific saccade-related effects in A1. **(A)** Average saccade onset-related delta ITC (top), theta (middle), and alpha (bottom) ITC within supragranular, granular, and infragranular cortical layers (coded by color). Color coded vertical dashed lines indicate the peak of the average layer-specific ITC. Peak delta ITC occurred at (supra, gran, infra): 300, 187, 212 ms. Peak theta ITC occurred at 271, 158, 118 ms and peak alpha ITC at 44, 54, 134 ms. Boxplots on the right show the average baseline ITC (base; from −300:300ms) and the average ITC 50 ms before and after the timing of the average layer-specific peak ITC indicated on the left across 62 experiments. Number of outliers excluded for visual purposes only (# supra/# gran/# infra) was theta: 0/0/1. Red brackets indicate significant differences based on Wilcoxon signed rank tests with Bonferroni correction. **(B)** Same organization as (A) for saccade offset-related ITC. Peak delta ITC occurred at (supra, gran, infra): 230, 142, 231 ms. Peak theta ITC occurred at 201, 45, 118 ms and peak alpha ITC at 177, 66, 0 ms.

In general, nonparametric tests were used to reduce the number of assumptions made about the data set being analyzed. Statistical comparisons of data presented in Figures 2 and 3 utilized Wilcoxon signed rank tests with Bonferroni correction correcting for frequency bands (3) and saccade events (2). Significance was determined to be p<0.008. To compare whether ITC was more strongly tied to saccade onset or offset for these data, Wilcoxon signed rank tests with Bonferroni correction were used. For data related to Figure 2, a correction for the number of frequencies (3) was used, which resulted in a significance level of p<0.017. For data related to Figure 3, a correction for the number of layers (3) and frequencies (3) was incorporated, which resulted in a significance level of p<0.006.

For an even more spatially fine-grained analysis, we detected significant ITC peaks across individual electrode channels (Figure 4). This method differed from that described above. For every channel and frequency, ITC values at each time point relative to saccade onset/offset were statistically evaluated using Rayleigh’s test of uniformity. The resulting p-values established the significance of phase alignment related to saccade onset/offset across cortical layers, frequencies, and time. Significant p values indicate a non-uniform distribution of phases. To correct for multiple comparisons, our p value was adjusted for the number of channels per recording (21), frequencies (41) and saccade events (2). After correction, significance was determined to be p<2.904×10^-5^. For each channel and frequency, local peaks in ITC from −500 to 500 ms around saccade events were determined using the “findpeaks” function in Matlab. Details about the ITC peaks occurring from −300 to 300 ms relative to saccade events were compiled. These details included information about ITC value, p value based on the Rayleigh test of uniformity, cortical depth, frequency, time, and normalized oscillatory amplitude at the same channel/frequency/time. Within each experiment, oscillatory amplitude was normalized across each channel and frequency by dividing by the mean amplitude from −300 to 300 ms around saccade onset and offset. From the pool of ITC peaks occurring between −300 and 300 ms, a subset of significant peaks (determined based on the Rayleigh p-value with Bonferroni correction described above) were further selected with their related details. This significant subset of ITC peaks is displayed in Figure 4A. Within this significant subset, comparisons between the timing, ITC value, and associated amplitude of the peaks related to saccade onset and offset are shown in Figures 4B-C. Statistical comparisons of the data presented in Figure 4C used Wilcoxon rank sum tests. A Bonferroni correction was applied for the number of frequencies (3) being tested. After correcting for these frequencies, significance was determined to be p<0.017. Significant differences in ITC timing were also evaluated using one sample t-tests. Figure 4D displays the cortical depth and timing of these significant peaks for each of the frequency bands of interest.

**Figure 4.**
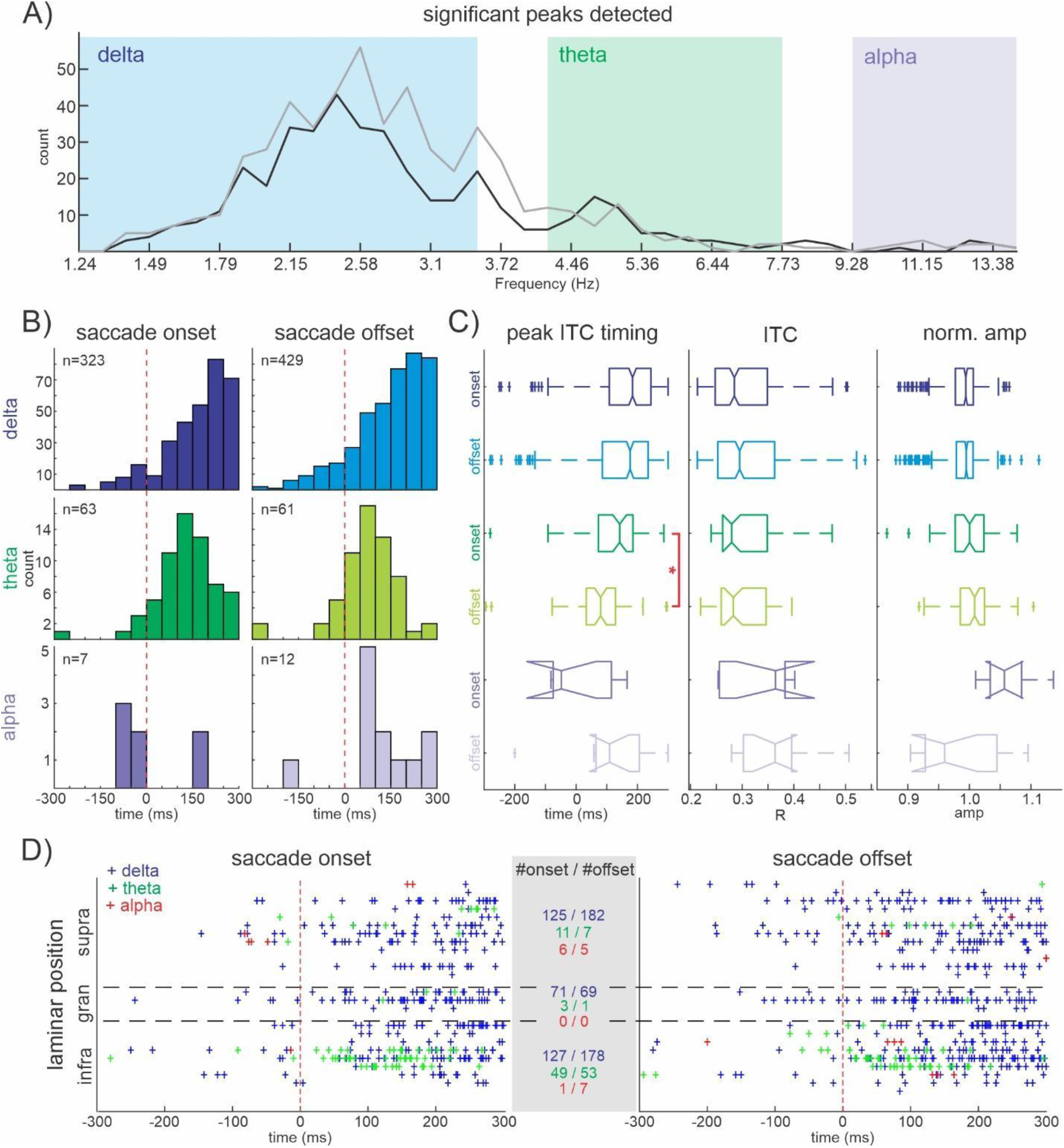
Saccade onset vs. offset related modulation of oscillatory activity in A1. (A) Distribution across frequencies of the 956 significant ITC peaks detected for saccade onset (black) and offset (grey) data. Blue/green/purple shaded boxes show the boundaries of the delta/theta/alpha bands of interest. **(B)** Histograms show the timing of the significant peaks relative to onset (left) and offset (right) for delta (top), theta (middle), and alpha (bottom) frequency bands. Vertical red dashed line at 0ms indicates the beginning of saccade onset and offset. **(C)** Boxplots describe the timing (left), ITC value (middle) and normalized oscillatory amplitude (right) of significant peaks detected in data across different frequency bands (coded by same colors as in (B)) aligned to saccade onset and offset. One onset and one offset outlier data point was excluded from the theta amplitude boxplots for visual purposes only. Red brackets with an asterisk denote a significant difference between onset and offset data based on Wilcoxon rank sum tests. Peak ITC timing onset vs offset: delta p=0.051, theta p=7.342×10-^4^, alpha p=0.099; ITC values onset vs offset: delta p=0.118, thetap=0.593, p=0.340; normalized amplitude onset vs offset: delta p=0.539, theta p=0.539, alpha p=0.017. **(D)** Laminar organization and timing of all significant ITC peaks for onset (left) and offset (right) data. Shaded box in the center lists the number of significant peaks for each frequency (by color) within each layer (# onset peaks/# offset peaks). Black horizontal dashed lines show the approximate border between layers. Vertical red dashed lines at 0ms indicate the beginning of saccade onset and offset.

## RESULTS

The goal of the current study was threefold. First, we aimed to examine whether eye movements provide a supramodal temporal context for the influx of environmental information in A1. Second, we wanted to determine if any eye movement related effects were layer specific, which would give an insight into possible circuit mechanisms. Finally, we aimed to disentangle whether any effects we found were related preferentially to saccade onsets vs. offsets. Addressing this last aim is significant given that there is a volley of visual information that enters the system following each saccade offset (i.e. at fixation onset) and that visual inputs influence auditory processing (Maddox et al., 2015; Atilgan et al., 2018). Therefore, effects more closely tied to saccade onset would indicate motor influences on auditory processing (a “motorsensory” interaction), while effects more closely related to saccade offset would indicate an effect related to visual input (multisensory interaction).

### Saccade-related modulation of CSD and MUA in A1

Neuroelectric activity was recorded in three macaques while they sat in a dark or dimly lit, sound attenuated recording chamber. Using linear array multielectrodes (Fig. 1A), field potential and multiunit activity (MUA) were concurrently recorded across all layers of A1. A total of 62 laminar A1 recordings were collected during 52 experimental sessions. Current source density (CSD) was calculated from the local field potential profiles (see Methods). Given the goal of the current study, examining CSD signals was advantageous over examining LFP signals as CSD reflects mostly local neuronal ensemble activity and is devoid of volume conduction (Kajikawa and Schroeder, 2011). Figure 1B shows representative CSD and MUA profiles in response to broadband noise burst (BBN) auditory stimuli. The CSD profile indexes the net direction of transmembrane current flow in the local neuronal ensemble across cortical layers, with sinks reflecting net inward current and sources reflecting net outward current (Mitzdorf, 1985; Schroeder et al., 1998). MUA profiles reflect neuronal ensemble firing rates across cortical layers. BBN responses in A1 displayed a robust modulation of both CSD and MUA, typical of a feedforward, driving input, with earliest onset and largest MUA response in layer 4 (Schroeder et al., 1998; Lakatos et al., 2007).

Eye position was recorded and saccade onsets and offsets were determined using the publicly available nonparametric method, Cluster Fix (König and Buffalo, 2014). Representative response profiles from the same recording location as Figure 1B to saccade onset and offset (Fig. 1C) exhibited a weaker modulation of CSD and MUA activity (compare Fig. 1C top with Fig. 1B). The slight decrease in MUA around saccade onset was similar to the saccade-related modulation of excitability in A1 previously reported by O’Connell and colleagues (2020). There is an influx of visual information that occurs near saccade offset/fixation, which occurs in dimly lit or dark environments. The small increase observed in saccade-related MUA could therefore be at least in part due to the multisensory influence of this visual information.

Using the Cluster Fix detection method, 6468 saccades were detected and manually verified. Median duration of the saccades was 43 ms and the median intersaccadic interval was 578 ms (Fig. 1D). The overall pace of the eye movements observed here was slightly slower than the typical 2-4 Hz that has been observed in previous NHP studies examining free viewing (Rajkai et al., 2008; Ito et al., 2011; Barczak et al., 2019). However, given that these NHPs were sitting in a darkened recording chamber without performing a task or receiving any rewards, it is reasonable to expect that the overall number and pace of eye movements made during each recording would be lower and slower than if they were actively engaged in a rewarding or engaging task.

### Saccade-related effects on oscillatory activity in A1

Previous studies of the visual system have shown that there is a cyclical modulation of excitability tied to the systematic patterns of fixations and saccades made by humans and NHPs while they actively sample the visual environment, which likely stems from oscillatory synchrony resulting from phase reset (Rajkai et al., 2008; Melloni et al., 2009; Barczak et al., 2019). Modulation of excitability related to saccade timing has also been demonstrated in the auditory system (O’Connell et al., 2020). To further inspect the mechanisms through which eye movements or related motor signals might provide a supramodal temporal context for the sampling of environmental information, we examined saccade-related modulation of oscillatory activity in A1. We first examined time-frequency intertrial coherence (ITC) derived from CSD epochs aligned to either saccade onset or offset. ITC was averaged across all channels spanning A1, from all 62 penetrations. Delta (1.2-3.5 Hz), theta (4.2-7.7 Hz), and alpha (9.3-14.2 Hz) frequency bands of interest were determined empirically by examining the boundaries of ITC increases in these grand averages (Fig. 2A). These empirically defined delta, theta, and alpha frequency bands closely matched those frequencies that have been traditionally studied when working with data related to NHPs (Lakatos et al., 2005; Haegens et al., 2011).

Figures 2B and 2C (left) show the frequency specific average ITC and oscillatory amplitude for all recordings aligned to saccade onset and offset. The peaks of the average delta and theta ITC related to saccade onset occurred slightly later (256 and 254 ms) than the peaks related to saccade offset (228 and 115 ms). While saccade durations varied (Figure 1D), most of the saccades examined in the current study had a duration less than 100 ms. Therefore, the earlier delta and theta ITC peaks related to saccade offset could, at least in part, be due to saccade onset and reflect a shift related to saccade duration. In contrast, peak alpha ITC related to saccade onset occurred earlier (44 ms) than that related to saccade offset (171 ms), which cannot be directly explained by a shift due to saccade duration.

To quantify these effects, peak ITC was statistically compared to a baseline ITC period for each experiment. As shown in Figure 2B (right), when compared to the base period, there was a significant post saccade event related increase in ITC within the delta and theta bands independent of whether epochs were aligned to saccade on or offset. The increase in ITC occurred without any significant modulation of oscillatory amplitudes within the same frequency bands (Fig. 2C, right). The lack of saccade-related amplitude change coupled with the significant ITC increase is indicative of a saccade-related modulation of ongoing delta and theta activity via phase reset and/or entrainment as opposed to a saccade-related evoked type response, which would have showed both a significant increase of oscillatory amplitude and ITC. While there was no significant modulation of alpha amplitude for either saccade onset or offset, there was a significant increase in alpha ITC related to saccade onset only. These results provide evidence for saccade-related modulation of A1 neuronal activity across multiple frequency bands.

To examine whether ITC was more strongly related to saccade onset or offset, average ITC related to each saccade event during the peak period was statistically compared across all experiments using Wilcoxon signed rank tests with Bonferroni correction. No significant differences in ITC during the peak periods were found between data related to saccade onset and offset (peak ITC: saccade onset vs offset; delta p=0.276, theta p=0.285, alpha p=0.249). Therefore, significant saccade-related modulation of ITC averaged across all cortical layers of A1 could not be statistically tied to either onset or offset events.

### Layer specific saccade-related effects in A1

To further evaluate the dynamics of saccade-related effects, we repeated the analyses described above for individual cortical layers. ITC data averaged within supragranular (supra), granular (gran), and infragranular (infra) channels across all A1 penetrations are shown in Figure 3 (left). Comparison of laminar ITC changes related to saccade onset and offset around the time of peak average ITC to a baseline period (Fig. 3, right) revealed that delta ITC after both saccade events was significantly larger across all experiments. This was true across all layers (Fig. 3A right; Wilcoxon signed rank tests with Bonferroni correction; saccade onset base vs peak: supra p=5.629×10^−5^, gran p=3.836×10^-4^, infra p=1.348×10^−4^; Fig. 3B, right; saccade offset base vs peak: supra p=2.111×10^−4^, gran p=9.010×10^−4^, infra p=1.837×10^−4^). There were no significant changes observed in delta oscillatory amplitude related to either saccade event across the same layers and experiments (data not shown; Wilcoxon signed rank tests with Bonferroni correction; saccade onset base vs peak: supra p=0.641, gran p=0.641, infra p=0.739; saccade offset base vs peak: supra p=0.891, gran p=0.777, infra p=0.335).

Across all experiments, theta ITC was also significantly larger after both saccade events (Fig. 3A right; Wilcoxon signed rank tests with Bonferroni correction; saccade onset base vs peak: supra p=4.266×10−4, gran p=0.002, infra p=0.004; Fig. 3B right; saccade offset base vs peak: supra p=4.380×10−4, gran p=0.005, infra p=1.689×10-4) while no changes were observed in theta oscillatory amplitude (data not shown; Wilcoxon signed rank tests with Bonferroni correction; saccade onset base vs peak: supra p=0.494, gran p=0.729, infra p=0.836; saccade offset base vs peak: supra p=0.353. gran p=0.282, infra p=0.554).

In contrast to the delta/theta frequencies, neither alpha ITC nor alpha oscillatory amplitude showed any saccade-related changes within individual layers (Fig. 3A right; ITC: Wilcoxon signed rank tests with Bonferroni correction; saccade onset base vs peak: supra p=0.035, gran p=0.112, infra p=0.196; Fig. 3B right; saccade offset base vs peak: supra p=0.201, gran p=0.646, infra p=0.198; Amplitude data not shown: saccade onset base vs peak: supra p=0.025, gran p=0.024, infra p=0.936; saccade offset base vs peak: supra p=0.060, gran p=0.216, infra p=0.229).

To evaluate whether observed laminar ITC effects were more strongly tied to either saccade onset or offset, layer and frequency specific saccade onset-related averages were compared to offset averages using Wilcoxon signed rank tests with Bonferroni correction. There were no significant differences between onset and offset-related peak ITC for any frequency or layer (saccade onset vs offset: delta: supra p=0.568, gran p=0.261, infra p=0.069; theta: supra p=0.468, gran p=0.402, infra p=0.022; alpha: supra p=0.563, gran p=0.592, infra p=0.540).

To summarize these results, significant increases in ITC without increases in oscillatory amplitude suggests that saccade related modulation of A1 neuronal ensemble activity across individual cortical layers is also characterized by the phase reset/entrainment of delta-theta oscillations. Despite our previous observation of a phase reset of alpha oscillations related to saccade onset when averaging activity across all layers, there were no saccade-related effects on alpha oscillations within individual cortical layers, likely indicating a weaker effect than that observed with lower frequencies. Directly comparing peak ITC related to saccade onset and offset did not yield any straightforward indication of whether the observed laminar effects were more closely tied to either onset or offset.

### Saccade onset vs. offset related modulation of oscillatory activity in A1

In a further attempt to disentangle effects tied to saccade onset (motor-related) and effects tied to saccade offset (visual input-related), we decided to implement a more fine-grained method for examining ITC peaks (see Methods). We determined significant ITC peaks within individual electrode channels (n=1117 total channels, median n=18 per recording), across all frequencies (n=41), and all A1 recording sites (n=62). In the time-period relative to saccade events (−300 to 300 ms), we detected 416 and 540 significant ITC peaks related to onset and offset, respectively. Figure 4A shows the frequency distribution of significant ITC peaks. Of these significant ITC peaks related to onset and offset, 393 and 502 occurred within our frequency bands of interest. This subset of 895 peaks occurred across 38 of 62 experiments (61.3%).

Within each frequency band, properties of the detected significant ITC peaks were compared. Histograms in Figure 4B show the timing of significant ITC peaks in different frequency bands related to saccade onset (left) and offset (right). Statistical evaluation of the timing of the peaks relative to a saccade event (Fig. 4C, left) using Bonferroni corrected

Wilcoxon rank sum tests showed that onset-related theta peaks occurred significantly later than those related to saccade offset (median timing: 142 ms vs 80 ms). This significant shift observed for theta peaks could be influenced by saccade duration. The median duration of saccades examined was around 43 ms (Fig. 1D) and therefore, the earlier timing of peaks related to saccade offset could reflect the peaks due to the non-aligned onset of those saccades. There were no significant differences between the timing of onset and offset related peaks in either the delta or alpha bands. Delta peaks related to saccade onset and offset occurred at 184 and 176 ms while alpha peaks occurred at −48 and 109 ms, respectively. Although it appears that there was a large difference between the timing of alpha peaks, the lack of significance could likely be due to the small sample size.

In addition to evaluating whether peak ITC timing related to saccade on- and offset were statistically different, we also evaluated whether peak ITC timing significantly differed from 0 ms. Results suggested that the timing of delta and theta peaks occurred significant later than both saccade onset and offset (One sample t-tests, delta: onset t(322)=27.62, p=6.148×10^-87^, offset t(428)=26.34, p=1.384×10^-91^; theta: t(62)=10.63, p=1.327×10^-15^, offset t(60)=5.76, p=3.098×10^-7^). Alpha ITC peaks related to saccade offset occurred significantly later than 0 ms while those related to saccade onset did not (One sample t-tests, alpha: onset t(6)=0.11, p=0.9160, offset t(11)=3.08, p=0.011).

To further examine saccade event-related differences between the significant peaks detected, we statistically evaluated the ITC value and normalized oscillatory amplitude of these peaks (Fig. 4C, middle and right). ITC values related to saccade onset and offset showed no significant differences across any frequency bands. There were also no significant differences between frequency specific oscillatory amplitudes related to saccade onset or offset. These null findings suggest that the current methods of detecting and evaluating peaks may not be best suited to determine if saccade-related effects are more strongly tied to either onset or offset (see Discussion).

Besides evaluating the frequency and timing of each significant ITC peak, our recording method also allowed us to approximate the cortical laminar location of these ITC peak events. Identifying the laminar location of these peaks is important because each peak indicates the significant modulation of A1 neuronal ensemble activity by saccade related events in the brain. This information can be used to further detail and localize saccade-related motor and visual effects on cortical activity. Figure 4D shows the timing and laminar organization of all significant ITC peaks relative to saccade onset (left) and offset (right) with frequency coded by color. The shaded box in the center lists the number of significant peaks within each frequency and cortical layer. While delta peaks occur in large numbers across all layers, theta and alpha peaks occurred more often in extragranular layers. Specifically, both saccade on- and offset-related theta ITC was most prevalent in the infragranular layers. In contrast, alpha band peaks were differentially localized: onset-related significant ITC peaks were detected mostly in the supragranular layers while those related to saccade offset were more evenly distributed between supragranular and infragranular layers.

## DISCUSSION

The goal of the current study was threefold. First, we aimed to examine whether saccades provide a supramodal temporal context for the influx of environmental information in A1. Second, determining if any saccade-related effects were layer specific in A1 would provide insight into possible mechanisms underlying the observed effects. Finally, we aimed to disentangle whether effects were specifically related to saccade on- or offset. Our approach revealed that oscillatory phase reset within delta and theta bands occurred at both saccade onset and offset. Although less consistent, we also found saccade-related phase reset of alpha oscillations. Further examination of these oscillatory effects across cortical layers of A1 showed that delta phase reset readily occurred across supragranular, granular and infragranular layers while theta and alpha effects occurred more often in extragranular layers. While our analyses did not yield any conclusive evidence that effects were more tied to either saccade on- or offset, our findings support the idea that saccade-related modulation of oscillatory activity occurs in A1 and confirms that saccade-related modulation of neuronal oscillations is supramodal. These conclusions underscore the importance of active vision in sensory modalities outside of vision and specifically highlight the relevance of eye position when examining auditory processing.

### Is saccade-related supramodal modulation in A1 more strongly tied to saccade onset (motor) or offset (visual) effects?

We observed saccade on- and offset-related increases of ITC across multiple frequency bands without an increase in oscillatory amplitude. These findings indicate that saccade-related effects are modulatory in nature and highlight the importance of motorsensory interactions – the influence of motor information on sensory processing. While data were collected during a “resting state” when NHPs were not directly involved in a task, the room was either dark or dimly lit. Given the lack of complete darkness along with the fact that visual information enters the system following each saccade offset (i.e. around fixation onset), and that visual inputs influence auditory processing (Maddox et al., 2015; Atilgan et al., 2018), our observed saccade-related effects in A1 could be due to the multisensory influence of the visual input at the end of each saccade, or alternatively due to motor-related signals.

We aimed to disentangle effects related to these two sensory and motor signals by evaluating whether our observed ITC effects were more strongly tied to saccade onset or offset. We hypothesized that effects more closely tied to saccade onset would indicate motor influences on auditory processing, while effects more closely related to saccade offset would indicate an effect of the visual input (multisensory interactions). Unfortunately, our statistical results did not provide a clear indication whether saccade-related effects were more strongly tied to onset or offset. This could be because the variation in saccade durations (Fig. 1D, median = 43 ms, std dev = 35.39 ms) was much smaller than any of the wavelengths of oscillations that are reset (wavelengths of 70-833 ms for 1.2-14.2 Hz frequencies examined). Therefore, our analysis might not be sensitive enough to pick up differences elicited by the shift associated with saccade duration. Studies conducted in complete darkness could be done to further elucidate this question. While we were not able to isolate whether the effects we observed were due to the motor component or to the incoming visual information, our results show that the rhythm at which the eye movements occur imposes physiological changes in A1 and emphasizes the importance of monitoring eye movements during tasks that utilize other sensory systems outside of visual.

### Potential anatomical origins of supramodal influence

If saccades provide supramodal context across multiple sensory cortices, how could the saccade-related effects anatomically reach these areas? Examination of saccade-related oscillatory effects across individual cortical layers showed a strong weighting of theta and alpha modulation in extragranular layers (Figure 4D). This laminar organization suggests the origin of effects may be outside of the direct lemniscal pathway from the ventral subdivision of MGB to A1. We discuss three subcortical structures that display properties and connectivity ideally suited for such multimodal modulation – the superior colliculus, pulvinar, and striatum. These three structures are thought to be involved in spatially guided behavior (Selemon & Goldman-Rakic 1988) and their ability to influence a vast number of cortical networks could support the type of supramodal modulation described in our results.

#### Superior Colliculus

The superior colliculus (SC) is heavily involved with different aspects of saccadic eye movements as well as with sensorimotor transformations occurring in different sensory modalities. Cells in the SC fire prior to the generation of a saccade (Munoz and Wurtz, 1995) and are involved in the initiation (Dorris et al., 1997) and execution (Krebs et al., 2010) of saccadic eye movements. The SC is also thought to be involved in sensorimotor transformations and contains visual, auditory, somatosensory and movement maps [reviewed in (Sparks, 1986; Gandhi and Katnani, 2011; Basso and May, 2017)]. Cells in the SC respond to visual, auditory and somatosensory stimuli individually and display additive, complex responses when multisensory stimuli are presented (Wallace et al., 1996). This structure’s involvement in more general action control and attention is further supported by the findings that activation occurs with other orienting movements such as pinnae, whisker, whole-body and head/limb movements (Basso and May, 2017).

Anatomical studies using lesions made in the deep layers of the SC revealed connections to the medial geniculus nucleus (MGN) and the inferior colliculus (Benevento and Fallon, 1975). Inputs from the inferior colliculus are relayed to MGN and then to auditory cortex (Pickles, 2015). Additionally, these lower layers of the SC receive inputs from somatosensory and auditory areas (May, 2006), which highlights the potential for a bidirectional transfer of information between SC and auditory cortex.

#### Pulvinar

The pulvinar is a large multisensory associative thalamic nucleus also with connections to multiple cortical areas [reviewed in (Shipp, 2003; Froesel et al., 2021)]. While the pulvinar’s extensive connections with different stages of visual processing (Robinson, 1993; Kaas and Lyon, 2007) has been widely explored, the pulvinar also has extensive connections to different cortical areas, sending and receiving inputs from other sensory modalities and higher order cortical areas. The ventral and dorsal portions of the pulvinar are differentially connected with early/late visual processing areas and parietal/frontal areas, respectively (Arcaro et al., 2015). Using neuroanatomical tracers, Cappe and colleagues (2009) showed that regions of the pulvinar had projections to parietal, auditory and premotor cortical areas and noted that the medial pulvinar was the site with the most overlap of different modalities. The medial pulvinar also has connections with prefrontal cortex (Romanski et al., 1997), the frontal eye fields (Trojanowski and Jacobson, 1974), parabelt (Pandya et al., 1994; Hackett et al., 1998; Scott et al., 2017), and areas of the temporal cortex involved with higher order auditory and language processing (Trojanowski and Jacobson, 1976, 1977).

The functional role of the pulvinar has not been clearly defined. Outside of its role in visual processing, Yirmiya & Hocherman (1987) showed that while some single units in the pulvinar responded to arm movements or auditory stimulation, other pulvinar units responded to both. Another study looking at single cell recordings in the pulvinar of NHPs showed that while some neurons responded to arm or eye movements independently, some cells had a larger response when these two movements occurred simultaneously, which the authors suggest, highlights the importance of motor intention in the pulvinar (Acuña et al., 1983). Recent studies have also described the pulvinar’s role in attention (Saalmann et al., 2012; Zhou et al., 2016; Fiebelkorn et al., 2019).

#### Striatum

The striatum receives input from the frontal eye field (Künzle and Akert, 1977), pulvinar (Parent et al., 1983; Lin et al., 1984; Kolomiets, 1993), MGB (Lin et al., 1984) as well as from different parts of the auditory cortex, but mostly those outside of A1 (Lin et al., 1984; Yeterian and Pandya, 1998). Like the pulvinar, many different functional roles have been attributed to the striatum. Hikosaka and colleagues (1993) suggests that the caudate facilitates saccades through signals sent from the caudate to the superior colliculus through the substantia nigra pars reticulata. The striatum may also contribute to the precision of movements using visual information from SC relayed through the pulvinar (Day-Brown et al., 2010). In addition to motor-related actions, the striatum has been shown to be involved in spatial processing (Karnath et al., 2002), sensory decision making (Romo et al., 1995; Guidotti et al., 2020), learning, and processing feedback (Cincotta and Seger, 2007; Behne et al., 2008). Interestingly, the striatum also seems sensitive to rhythmic information. Bengtsson & Ullen (2006) showed that the striatum is involved in processing melodic and rhythmic information while humans played piano. Araneda and colleagues (2017) showed that there is a supramodal neural network for beat processing that involves the putamen. Taken together, these results highlight a potential role for the striatum to track and relay (quasi-) rhythmic motor-related information to other corticothalamic networks to enhance movements.

### Importance and implications of auditory motor connections

Since primate ears do not move as robustly as eyes, active sensing in the auditory cortex is not as straightforward as in the visual or somatosensory systems where movements (e.g. eyes and hands) are actively utilized to obtain modality-specific inputs. Saccade-related supramodal modulation suggests that the motor system could lead to coherence of activity across multiple sensory systems and influence sensory processing on a large scale. In addition to the current results showing that saccades can reset the phase of ongoing oscillations in A1, saccades have also been shown to modulate excitability in A1 (O’Connell et al., 2020). Highlighting the fact that these kinds of motor signals can impact sensory processing at the very earliest stages, saccadic eye movements have been shown to lead to oscillatory-type movement of the eardrum even when no auditory stimuli are presented (Gruters et al., 2018). These motorsensory influences are not unidirectional in nature. Auditory stimuli have also been shown to have effects on motor related processes as well. The presentation of auditory stimuli can lead to transient changes in pupil size (Wang et al., 2014) and eye movements tend to stop prior to predictable auditory targets (Abeles et al., 2020). In addition to reacting to salient stimuli, the motor system can also respond to intricate changes in perception. For example, when complex auditory patterns emerged amid a continuous auditory stimulus sequence with consistent presentation rates, pupil diameter could differentiate and track these patterns despite there being no physical break in the timing between individual stimuli (Barczak et al., 2018).

The motor cortex’s involvement in more complex cognitive processes like sensory-motor transformations, action-related information processing, attention, and prediction [reviewed by (Rizzolatti and Luppino, 2001; Schubotz, 2007; Morillon et al., 2015)] further emphasizes the importance of these bidirectional interactions of sensory and motor systems. Using fMRI, Archakov and colleagues (2020) showed that when NHPs listen to sounds they previously learned to produce using a “monkey piano” they show activation of hand and arm representations of the motor cortex and the putamen. Incorporating rhythmic movement routines led to enhanced attentional processing (Schmidt-Kassow et al., 2013) and increased perception of relevant stimuli while suppressing irrelevant ones (Morillon et al., 2014). Since ongoing neuronal oscillations reflect alternating states of excitability within a neuronal ensemble (Buzsáki, 2006) and modulation of these inherent rhythms has been proposed as a mechanism by which processes like attentional selection could occur (Rajkai et al., 2008; Lakatos et al., 2013), it has been proposed that rhythmic or quasi-rhythmic motor actions (i.e. saccades) might use these oscillatory fluctuations to facilitate more complex cognitive processes (Melloni et al., 2009; Morillon and Schroeder, 2015).

## Acknowledgements

Support for this work was provided by NIH grants R01DC012947, P50MH109429, and R01MH109289.

